# Alpha/beta power decreases during episodic memory formation predict the magnitude of alpha/beta power decreases during subsequent retrieval

**DOI:** 10.1101/2020.07.08.193763

**Authors:** Benjamin J. Griffiths, María Carmen Martín-Buro, Bernhard P. Staresina, Simon Hanslmayr, Tobias Staudigl

## Abstract

Episodic memory retrieval is characterised by the vivid reinstatement of information about a personally-experienced event. Growing evidence suggests that the reinstatement of such information is supported by reductions in the spectral power of alpha/beta activity. Given that the amount of information that can be recalled depends on the amount of information that was originally encoded, information-based accounts of alpha/beta activity would suggest that retrieval-related alpha/beta power decreases similarly depend upon decreases in alpha/beta power during encoding. To test this hypothesis, seventeen human participants completed a sequence-learning task while undergoing concurrent MEG recordings. Regression-based analyses were then used to estimate how alpha/beta power decreases during encoding predicted alpha/beta power decreases during retrieval, on a trial-by-trial basis. When subjecting these parameter estimates to group-level analysis, we find evidence to suggest that retrieval-related alpha/beta (7-15Hz) power decreases fluctuate as a function of encoding-related alpha/beta power decreases. These results suggest that retrieval-related alpha/beta power decreases are contingent on the decrease in alpha/beta power that arose during encoding. Subsequent analysis uncovered no evidence to suggest that these alpha/beta power decreases reflect stimulus identity, indicating that the contingency between encoding- and retrieval-related alpha/beta power reflects the reinstatement of a neurophysiological operation, rather than neural representation, during episodic memory retrieval.

## Introduction

Episodic memory refers to detail-rich, durable memories of personally-experienced events (Tulving, 2002). During the formation of these memories, large quantities of information about the event need to be represented and encoded by the cortex, while the later retrieval of these memories similarly hinges upon the representation of this previously-encoded information within the cortex. As such, an intuitive contingency exists in which the amount of information encoded dictates the amount of information that can later be retrieved (Tulving & Thomson, 1973). However, it is unclear whether such a relationship is observable on the neural level. Do the neurophysiological phenomena that facilitate information representation during episodic memory formation predict the magnitude of neurophysiological phenomena that facilitate information representation during subsequent retrieval? Here, we examine this idea.

To successfully encode and retrieve details about an episodic memory, the neural signal representing these details needs to be elevated above the noise generated by ongoing, task-irrelevant neuronal activity (Harris & Thiele, 2011). One potent form of background noise comes from “noise correlations” – that is, the synchronised firing of neurons unrelated to the signal of interest. When task-irrelevant neurons synchronise their firing, their summed activity conceals the comparatively small activity generated by the signal of interest. Therefore, the desynchronisation of these task-irrelevant neurons can enhance information representation within the neocortex by reducing background noise (Hanslmayr et al., 2012, 2016). Given that these noise correlations share a strong positive correlation with local field potential (LFP; Cui et al., 2016), one could speculate that the ubiquitous reduction in alpha/beta power that arises during task engagement (e.g., Crone et al., 1998; Krause et al., 1994; Pfurtscheller et al., 1996) may reflect a reduction in underlying noise correlations. In line with this idea, growing evidence suggests that alpha/beta power decreases support the representation of information encoded within, and retrieved from, episodic memories (Griffiths et al., 2020; Griffiths, Mayhew, et al., 2019; Karlsson et al., 2020; Martín-Buro et al., 2020). For example, a simultaneous EEG-fMRI study demonstrated that the magnitude of alpha/beta power decreases directly correlated with the amount of stimulus-specific information represented within the BOLD signal during both perception and episodic memory retrieval (Griffiths, Mayhew, et al., 2019). Notably, these alpha/beta power decreases did not represent stimulus-specific information, but rather provided the conditions which benefit information representation (for complementary evidence, see Weisz et al., 2020). One could therefore hypothesise that reductions in alpha/beta power serve to reduce noise rather than boost signal. Taking these findings together, it appears that decreases in alpha/beta power reflect a neurophysiological phenomena that provides beneficial conditions for information representation within the cortex (Hanslmayr et al., 2012).

Given that information represented within the cortex during retrieval is contingent on the information originally encoded (Tulving & Thomson, 1973), we hypothesised that retrieval-related alpha/beta power decreases are similarly contingent on encoding-related alpha/beta power decreases. To test this, seventeen participants completed a sequence learning task while undergoing concurrent MEG recordings (see figure 1a). We then modelled alpha/beta power during retrieval as a function of alpha/beta power during initial encoding (see figure 1b). In line with our hypothesis, we found evidence to suggest that, on a trial-by-trial basis, alpha/beta power decreases during episodic memory retrieval can be predicted by the associated alpha/beta power decreases observed during encoding.

**Figure 1.**
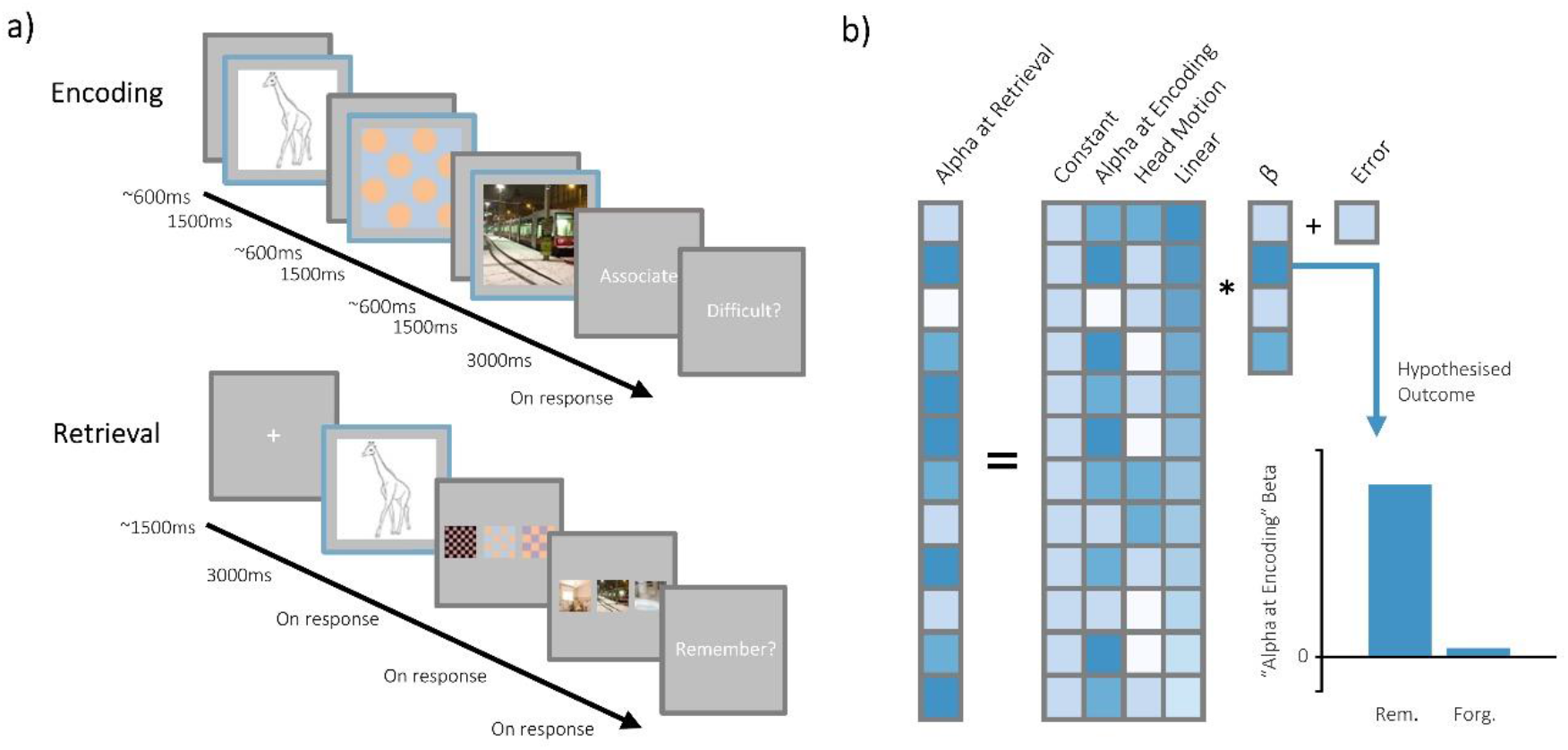
Overview of behavioural task and analytical approach. **(a)** Paradigm schematic. Participants were presented with a sequence of three visual stimuli. The sequence began with a line drawing of an object, followed by a feature and a scene (each with a brief fixation cross shown between). Participants were then given a short interval to create a mental image incorporating the three stimuli, and asked to rate how difficult it was to create the association. After a distractor task, participants were presented with the object as a cue and asked to recall the pattern and the scene. After selection, participants had to rate how confident they felt about their response. Windows outlined in blue depicted the epochs analysed. **(b)** For every participant, a multiple regression model was created in which alpha/beta power on a given retrieval trial was predicted based on alpha/beta power at encoding, head motion, linear drift, and a constant. The parameter estimate describing the linear relationship between alpha/beta power at encoding and alpha/beta power at retrieval was extracted for every sensor-frequency pair of every participant, and these estimates were subjected to inferential statistical analysis. We hypothesised that a positive linear relationship would be observed across participants, indicating that the alpha/beta power during encoding predicts alpha/beta power during retrieval. Furthermore, we hypothesised that this effect would be restricted to successfully recalled items, as unsuccessful retrieval does not elicit meaningful memory-related fluctuations in alpha/beta power.

## Methods

### Data statement

The data analysed here is taken from a previous dataset (Griffiths et al., 2020). Exclusion criteria match those that were pre-registered for the previous study.

### Participants

Seventeen participants were included in the final analysis (mean age = 24.9; age range = 20-32; 11 female, 6 male; 14 right-handed, 3 left-handed). These participants received course credit or financial reimbursement in return for their participation. An additional eleven participants were excluded (n=1 for excessive head movement, n=4 for poor data quality, n=6 for having fewer than 15 forgotten triads in the pre-processed data). These exclusion criteria matched those reported by Griffiths and colleagues (2020). Ethical approval was granted by the Research Ethics Committee at the University of Birmingham (ERN_15-0335), complying with the Declaration of Helsinki.

### Experimental design

Each participant completed a visual associative memory task (see figure 1a). During encoding, participants were presented with a line drawing of an object (either “animate” or “inanimate”; 50% of trials belonged to each category, evenly distributed throughout the experiment), a feature (“polka-dot” or “chequered”), and a scene (“indoor” or “outdoor”). Each stimulus was presented for 1500ms, with a jittered 600ms (±100ms) fixation cross presented between each stimulus. Participants were then given a short interval (3000ms) to create a mental image incorporating the three stimuli to help them recall the stimuli during a later memory test. After associating 48 triads, participants completed a short visual perceptual distractor task. A retrieval task then followed the distractor. Here, participants were presented with the line drawing (for 3000ms) and asked to recall the mental image they created during the encoding phase. Then, participants were presented with three patterns (one correct and two lures) and asked to identify the pattern associated with the line drawing. After responding, this process was repeated for the scene stimuli, again using the correct stimulus and two lures. Participants were asked to recall all 48 triads learnt in the earlier encoding phase. Participants completed four blocks of this task (192 trials in total). For all responses, participants used two non-magnetic, single-finger optical response pads. The left pad allowed participants to cycle through the possible responses, and the right pad allowed participants to confirm their selection.

### MEG acquisition

MEG data was recorded using a 306-channel (204 gradiometers, 102 magnetometers) whole brain Elekta Neuromag TRIUX system (Elekta, Stockholm, Sweden) in a magnetically shielded room. Participants were placed in the supine position for the duration of the experiment. The MEG was continuously recorded at a sampling rate of 1000Hz. The head shape of each participant (including nasion and left/right ear canal) was digitised prior to commencing the experiment. Continuous head position indicators (cHPI) were recorded throughout. The frequencies emitted by the cHPI coils were 293Hz, 307Hz, 314Hz and 321Hz. Magnetometer data was excluded from the main analysis as they contained substantial noise that could not be effectively removed or attenuated.

### MEG preprocessing

All data analysis was conducted in Matlab using Fieldtrip (Oostenveld et al., 2011) and custom scripts. First, the data was lowpass filtered at 165Hz to remove the signal generated by the HPI coils. Second, the data was epoched around each event of interest. At encoding, the epochs reflected the time windows where each stimulus was presented. At retrieval, the epochs reflected the time window when the object cue was presented. Encoding epochs began 2000ms before stimulus onset and ended 3500ms after onset (that is, 2000ms after stimulus offset). Retrieval epochs began 2000ms before stimulus onset and ended 4500ms after onset (that is, 2000ms after stimulus offset). Third, independent components analysis was conducted, and any identifiable eye-blink or cardiac components were removed. Fourth, the data was visually inspected, and any artefactual epochs or sensors were removed from the dataset (mean percentage of trials removed: 18.0%; range: 5.7-32.2%).

### Movement correction

To help attenuate motion-related confounds in the spectral power analyses, a trial-by-trial estimate of motion was calculated. First, the data was highpass filtered at 250Hz. Second, the data was epoched into trials matching those outlined in the section above. Third, the envelope of the signal in each epoch was calculated (to avoid issues of mean phase angle difference in cHPI signal across trials). Fourth, the envelope was averaged over time to provide a single value for each epoch and channel. Fifth, the dot product was computed across sensors between the first epoch and every other epoch (algebraically: 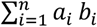, where *n* is the number of channels, ***α_i_*** is the power at sensor ***i*** during the first trial, and ***b_i_*** is the power at sensor ***i*** during the trial of interest). This provided a single value (between zero and infinity) for each trial that described how similar the topography of that trial was to the first trial – the higher the value, the more similar the topographies are between the two trials (with the assumption that the more dissimilar a topography is to the starting topography, the more the head has deviated from its starting position). These values were entered as a regressor of no interest in the central multiple regression analyses.

### Time-frequency decomposition

Sensor-level time-frequency decomposition was conducted on the two epochs (encoding and retrieval). The preprocessed data was first convolved with a 6-cycle wavelet (0 to 1.5 seconds for encoding; 0 to 3 seconds for retrieval, in steps of 50ms; 3 to 40Hz; in steps of 0.5Hz). Second, planar gradiometers were combined by summing the power of the vertical and horizontal components. Third, for encoding epochs only, power was then averaged over the three stimulus presentation windows of each triad to provide mean power during the encoding of the triad. Any triads where one or more epochs had been rejected during preprocessing were excluded at this stage. We averaged spectral power across the three windows as we reasoned that this approach would be most sensitive to changes in spectral power that predicted the quantity of information later recalled. Fourth, post-stimulus power for the encoding and retrieval epochs were averaged across time. We did this as we had no expectation of when alpha/beta power during encoding would overlap with alpha/beta power during retrieval.

### Multiple regression analysis

Our central analyses were inspired by the multi-level approach frequently used in fMRI research, where participant-level effects are first estimated using general linear models, and the resulting parameter estimates are subjected to group-level inferential statistics (see figure 1b). To this end, three first-level linear regression models were run for each participant: one model described how retrieval success related to spectral power at retrieval (the “memory performance” model), one described how encoding spectral power related to retrieval spectral power during successful memory retrieval, and one model described how spectral power during encoding related to spectral power during unsuccessful memory retrieval (the “spectral power” models). We used three models (rather than a single model incorporating individual regressors for both memory performance and spectral power at encoding) as we would anticipate that there would be significant collinearity between memory performance and spectral power at encoding (i.e. the subsequent memory effect; for reviews, see Hanslmayr & Staudigl, 2014; Kim, 2011). By using separate models for memory performance and spectral power at encoding, we avoid potentially spurious model fits driven by collinearity introduced by the subsequent memory effect.

For the first linear model, three regressors and a constant were used to predict spectral power during the retrieval epoch. The first regressor described whether the triad was successfully recalled or not. If the triad was recalled in its entirety, a value of ‘1’ was used. If none of the triad was recalled, a value of ‘0’ was used. The second regressor described motion at retrieval, and served to supress variance introduced by this potential confound. The third regressor reflected a linear drift, and served to supress variance introduced by signal autocorrelation over time. The outcome variable was spectral power at retrieval for a given channel-frequency pairing. The model was fitted for every channel-frequency pairing individually. Each fitting provided a parameter estimate describing how the binary memory regressor predicted spectral power during retrieval. This parameter estimate was then standardised by dividing the standard error of the fit, bringing every parameter estimate into the same unit space and facilitating across-participant comparisons.

For the second and third linear models, the data was restricted to successfully retrieved and forgotten triads respectively. Five regressors and a constant were used to predict spectral power during the retrieval epoch. The first regressor described alpha/beta power at encoding. The second and third regressors described motion at encoding and retrieval, and served to supress variance introduced by these potential confounds. The fifth and sixth regressors reflected trial numbers at encoding and retrieval, and served to supress variance introduced by signal autocorrelation over time. The outcome variable matched that of the first model, and model estimation was conducted in the same manner as reported above.

### Statistical analysis

For statistical analysis of the memory performance model, the standardised parameter estimates were contrasted against the fit expected by chance. This chance-level fit was estimated by randomly permuting the values of the outcome vector relative to the predictor matrix 1000 times, and then calculating the mean of these chance-level fits. The t-values for each participant were contrasted against the chance-level fits in a dependent-samples, cluster-based permutation test (2000 permutations, alpha threshold = 0.05, cluster alpha threshold = 0.05, minimum neighbourhood size = 3; Maris & Oostenveld, 2007). Note that replacing the permuted chance null hypothesis with an absolute zero null hypothesis produces synonymous results.

For statistical analysis of both spectral power models, a region of interest was defined based on the cluster uncovered when analysing the memory performance model. This helped focus our analysis on spectral power related to episodic memory processes, and not to other co-occurring cognitive phenomena such as perception or attention. The parameter estimates of each participant were averaged across all channel-frequency pairings included in the cluster, leaving a single average parameter estimate for each participant and each model. For the initial analysis, these parameter estimates were contrasted against the chance-level fit in a dependent samples t-test (for successfully recalled and forgotten models separately). Then, the parameter estimates of the two models were directly contrasted in permutation-based, dependent samples t-test.

One may sense that this latter analysis is circular – the contrast of parameter estimates for hits and parameter estimates for misses is conducted using a region of interest derived from a previous analysis of hits and misses. However, this is not the case. The original contrast of the memory performance model approximates the “retrieval success effect”, where the mean spectral power for hits is contrasted with the mean spectral power for misses. As such, one can view the cluster as being those features sensitive to differences in mean spectral power between conditions. When we subsequently build the spectral power regression models for hits and misses separately, the mean spectral power for each memory condition is captured by the constant included in their respective models. With the mean spectral power captured in the constant (and, therefore, explicitly *not* in the parameter estimates of the specified regressors), the contrast of the parameter estimates between hits and misses can be viewed as an orthogonal contrast to the “retrieval success” contrast that was conducted first.

### Time-generalisation analysis

To understand the temporal dynamics of the central effect, we computed a time-generalisation matrix to explore how alpha/beta power at every time point during encoding correlated with alpha/beta power at every time point during retrieval. Here, the analysis matched that of the multiple regression approach described above with one key exception: power was not averaged over time. Instead, the linear models were computed for every channel x frequency x time-at-encoding x time-at-retrieval combination (restricted to channels and frequencies included within the memory-related cluster). The derived standardised parameter estimates were then averaged over channels and frequencies to provide a two-dimensional matrix describing how alpha/beta power at encoding correlated with alpha/beta power at retrieval for every time point during encoding (dimension 1) and every time point during retrieval (dimension 2). As before, this was done separately for remembered and forgotten trials.

To identify time windows in which a substantial correlation between encoding and retrieval alpha/beta power exists, the time generalisation matrices were contrasted against the fits expected by chance (calculated as in the previous section) in a dependent-samples, cluster-based permutation test (2000 permutations, alpha threshold = 0.05, cluster alpha threshold = 0.05, Maris & Oostenveld, 2007). This was done for remembered and forgotten trials separately. Another dependent-samples, cluster-based permutation test then assessed the difference in fits between remembered and forgotten trials.

### Linear discriminant analysis

To examine whether alpha/beta power contained stimulus-specific information, linear discriminant analysis (LDA) was conducted on the time-series data of the encoding epochs. Specifically, LDA was used to identify whether the stimulus subcategories could be distinguished from one another [i.e. animate vs. inanimate subcategories for the object stimuli; polka-dot vs. chequered subcategories for the feature stimuli; indoor vs. outdoor subcategories for the scene stimuli]. LDA analysis was conducted using the MPVA light toolbox (Treder, 2020). LDA was first run on broadband amplitude to test whether the electrophysiological signal in its entirety contained stimulus-specific information. For ease of reading, we describe the process for comparing “animate vs. inanimate” object stimuli, however the pipeline generalises to that used for the “polka-dot vs. chequered” and “indoor vs. outdoor” contrast. First, the dimensionality of the data was reduced. Sensor-level data was converted into components using Principle Component Analysis (PCA) and the 50 components that explained the most variance in the data were selected. We elected to take 50 components as this provided a reasonable trade-off between having sufficient components to detect differences and having a sufficient ratio of trials to components to avoid over-fitting (approximately three trials to one feature). Second, the trials were split based on their subcategory (i.e. “animate” or “inanimate”). Third, trial numbers between the two subcategories were balanced by taking all trials from the subcategory with a smaller number of instances, and then, for every one of these trials, selecting the trial of the opposite category that occurred nearest in time to this trial. This approach minimised differences between the subcategories that could be ascribed to signal autocorrelation. Fourth, the data was randomly partitioned into five folds (with an equal number of trials from both categories included within a fold). Fifth, weights that best separated the two subcategories were determined based on data in four of these folds (i.e. the training set), and then applied to the fifth fold (i.e. the testing set). This process was repeated in a cross-validated manner, using each fold as the testing set in turn. This process was conducted for every sample point separately. Classification performance was calculated by assigning each test trial a binary value (1 for when the predicted label matched the observed label; 0 for when the labels did not match) and then averaging across trials to provide a value between 0 and 1 describing the percentage of times the classifier produces the correct outcome. The accuracy of the classifier for each participant were pooled and entered into a group-level within-subject t-test, where they were contrasted against chance (estimated by permuting test labels relative to the test data across trials and deriving the associating accuracy) in a dependent-samples, cluster-based permutation test (using parameters as before).

To test whether alpha/beta power contained stimulus-specific information, this analysis pipeline was re-run from the beginning using alpha/beta power in place of broadband amplitude. Alpha/beta power was estimated by taking the absolute of the envelope of the narrowband filtered signal (IIR filter: 7-15Hz, ensuring that all frequencies included in the main cluster effect were included in this analysis).

## Results

### Alpha/beta power decreases during retrieval correlate with memory performance

In the first instance, we aimed to verify the presence of a memory-related reduction in alpha/beta power during successful memory retrieval. To this end, spectral power during the retrieval epoch was modelled as a linear combination of memory performance (entire sequence recalled vs. none of the sequence recalled), head motion, and trial number for every participant. The resulting parameter estimated describing how memory performance predicted spectral power was then extracted for each participant, and a group-level contrast was conducted to see whether these coefficients consistently deviated from chance across participants. Here, the successful retrieval of a triad was accompanied by a decrease in alpha/beta power relative to forgotten triads [p_clus_ = 0.022, summed t-statistic = −931.24, cluster size = 345, Cohen’s dz = 0.65]. The reported cluster was observed over occipital and left parietal cortices, and had a frequency range of 7-15.5Hz (see figure 2a and 2d). This matches numerous previous studies linking alpha/beta power decreases to successful memory retrieval (e.g. Griffiths et al., 2020; Griffiths, Mayhew, et al., 2019; Karlsson et al., 2020; Khader & Rösler, 2011; Martín-Buro et al., 2020; Michelmann et al., 2016; Waldhauser et al., 2016).

**Figure 2.**
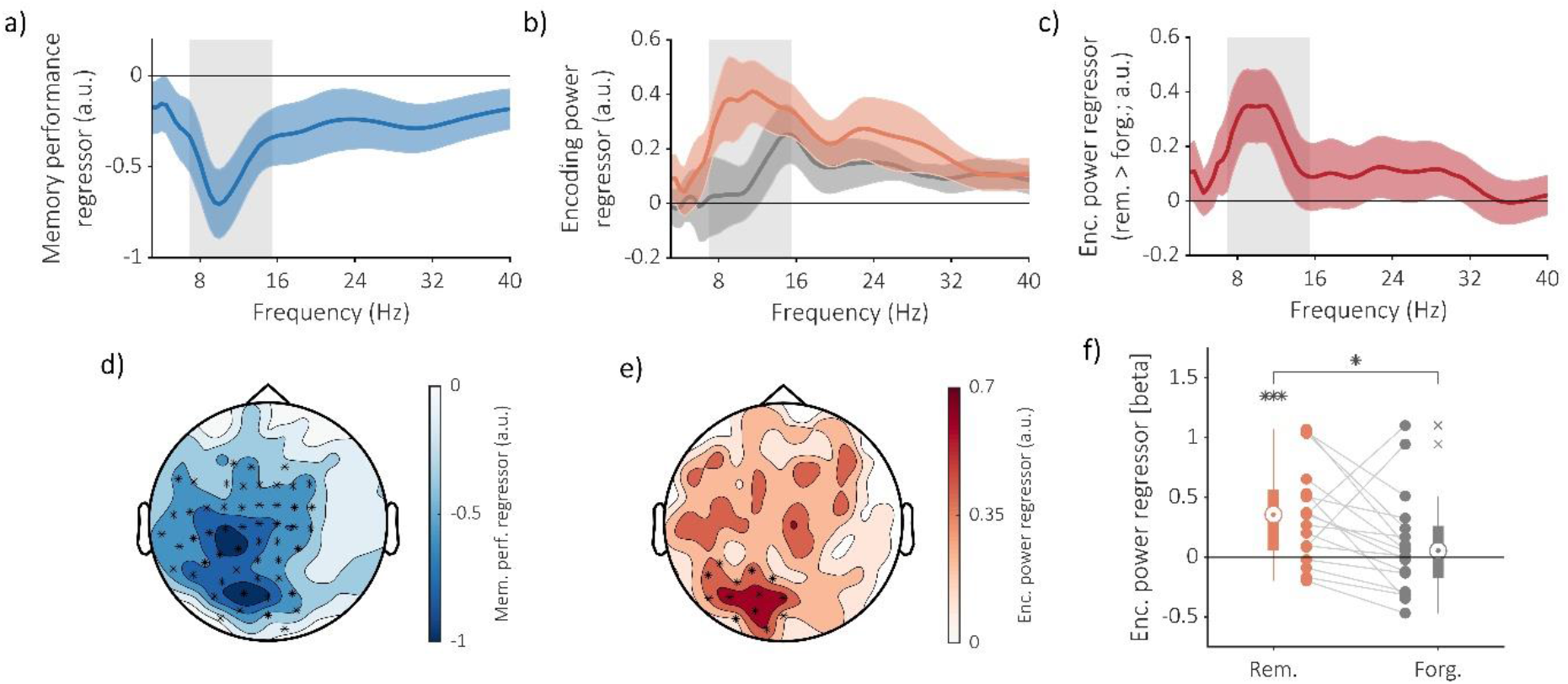
Retrieval-related alpha/beta power decreases can be predicted based on encoding. **(a)** Frequency spectrum depicting difference in power as a function of memory performance. Alpha/beta power exhibits a significant decrease during successful sequence retrieval relative to unsuccessful retrieval. Dark line indicates mean standardised parameter estimate (averaged across participants and sensors included in the significant cluster); shaded error bar depicts standard error of standardized parameter estimate across participants; grey shaded region depicts frequencies included in the significant cluster. **(b)** Frequency spectrum depicting how encoding power predicts retrieval power for hits (in orange) and misses (in grey). Encoding power within the alpha/beta band significantly predicts retrieval power for hits, but not misses. Dark line indicates mean standardised parameter estimate (averaged across participants and sensors included in the region of interest); shaded error bar depicts standard error of standardised parameter estimate across participants; grey shaded region depicts frequencies included in the region of interest. **(c)** Frequency spectrum depicting memory-related differences in how encoding power predicts retrieval power. Encoding power within the alpha/beta band predicts retrieval power to a significantly greater degree than misses. Dark line indicates mean standardised parameter estimate (averaged across participants and sensors included in the region of interest); shaded error bar depicts standard error of standardised parameter estimate across participants; grey shaded region depicts frequencies included in the region of interest. **(d)** Topography depicting difference in power (averaged across participants and frequencies included in significant cluster) as a function of memory performance. Alpha/beta power decreases are most prominent over left occipital and parietal sensors. Crosses indicate electrodes included in cluster when comparing the observed memory parameter estimate to chance. **(e)** Topography depicting the predictability of retrieval alpha/beta power (averaged across participants and frequencies included in region of interest) for remembered sequences. Predictability is widespread, with the greatest correlation over the same occipital regions observed in the memory-related contrast (see panel d). Crosses indicate electrodes included in unrestricted cluster analysis comparing the correlation between encoding and retrieval power for remembered sequences to chance. **(f)** Boxplot depicting the predictability of retrieval alpha/beta power based on encoding power, as function of memory performance. Across participants (as individual dots), remembered items showed a significant relation, whereas misses did not.

### Encoding-related alpha/beta power decreases predict subsequent retrieval-related alpha/beta power decreases

We then addressed our key hypothesis: is the retrieval-related decrease in alpha/beta power dependent on the alpha/beta power decrease during encoding? To this end, spectral power during the retrieval epoch was modelled as a linear combination of spectral power during the encoding epoch [within the cluster uncovered when analysing the memory performance model], head motion, and trial number. The resulting standardised parameter estimates for every channel-frequency pairing within the memory-related cluster were averaged to provide a single value describing the correlation between encoding- and retrieval-related spectral power for remembered items (for each participant separately). These values were entered into a group-level contrast to see whether these parameter estimates consistently deviated from chance across participants. Here, we observed a significant positive correlation between encoding and retrieval alpha/beta power on a trial-by-trial level [t(16) = 3.16, p < 0.001, Cohen’s d_z_ = 0.76; see figure 2b (hits in red, misses in grey)], indicating that fluctuations in alpha/beta power during successful retrieval can be predicted by fluctuations in alpha/beta power during the initial encoding. Unrestricted cluster-based analysis produced synonymous results: a significant positive correlation was observed between encoding and retrieval alpha/beta power on a trial-by-trial level [p_clus_ = 0.034, summed t-statistic = 156.08, cluster size = 62, Cohen’s d_z_ = 0.63]. This cluster was present over the occipital cortex, and had a frequency range of 9-14.5Hz (see figure 2e for topography), complimenting the preceding finding by demonstrating that retrieval-related alpha/beta power decreases can be predicted by prior encoding-related power decreases.

We then repeated this analysis on the forgotten items. Here, no correlation was found between encoding and retrieval alpha/beta power [t(16) = 0.87, p = 0.198, d = 0.21; see figure 2b], suggesting that this effect is unique to when information is successfully encoded and subsequently recalled.

We then formally tested the differences in fits between hits and misses. Here, we found evidence to suggest that the fit for recalled items is significantly stronger than the fit for forgotten items [t(16) =1.99, p = 0.032, d = 0.48; see figure 2c and 2f], further supporting the idea that this effect is a memory-related phenomena.

### Temporally-brief encoding-related alpha/beta power decreases predict temporally-extend retrieval-related power decreases

To examine the temporal specificity of these effects, a time-generalisation matrix was computed in which alpha/beta power for every time-point at encoding was correlated with every time-point at retrieval using the same linear model-based approach as above (this time, averaged over channels and frequencies within the memory-related cluster). For remembered items, two clusters were identified. The first cluster uncovered a significant correlation between encoding- and retrieval-related alpha/beta power, which included encoding time-points from 100ms to 1500ms and retrieval time-points ranging from 125ms to 1950ms [p_clus_ = 0.006, summed t-statistic = 3484.52, cluster size = 1425, Cohen’s d_z_ = 0.59]. While temporally broad, inspection of the time generalisation matrix suggests that this cluster peaked at encoding time-points between 500 and 1000ms and at retrieval time-points between 300 and 1800ms (see figure 3a). The second cluster uncovered a significant correlation between encoding- and retrieval-related alpha/beta power, which included encoding time-points from 400ms to 1500ms and retrieval time-points ranging from 2375ms to 3000ms [p_clus_ = 0.021, summed t-statistic = 2398.98, cluster size = 892, Cohen’s d_z_ = 0.65]. These results indicate that the correlated alpha/beta power decreases have different time-courses during encoding and retrieval. Indeed, the timing of the encoding-related power decrease matches that of the subsequent memory effect (e.g. Fell et al., 2008; Griffiths et al., 2016; Hanslmayr et al., 2009), while the timing of the retrieval-related power decreases matches that of the retrieval success effect (e.g. Karlsson et al., 2020; Martín-Buro et al., 2020; Michelmann et al., 2016; Waldhauser et al., 2016).

**Figure 3.**
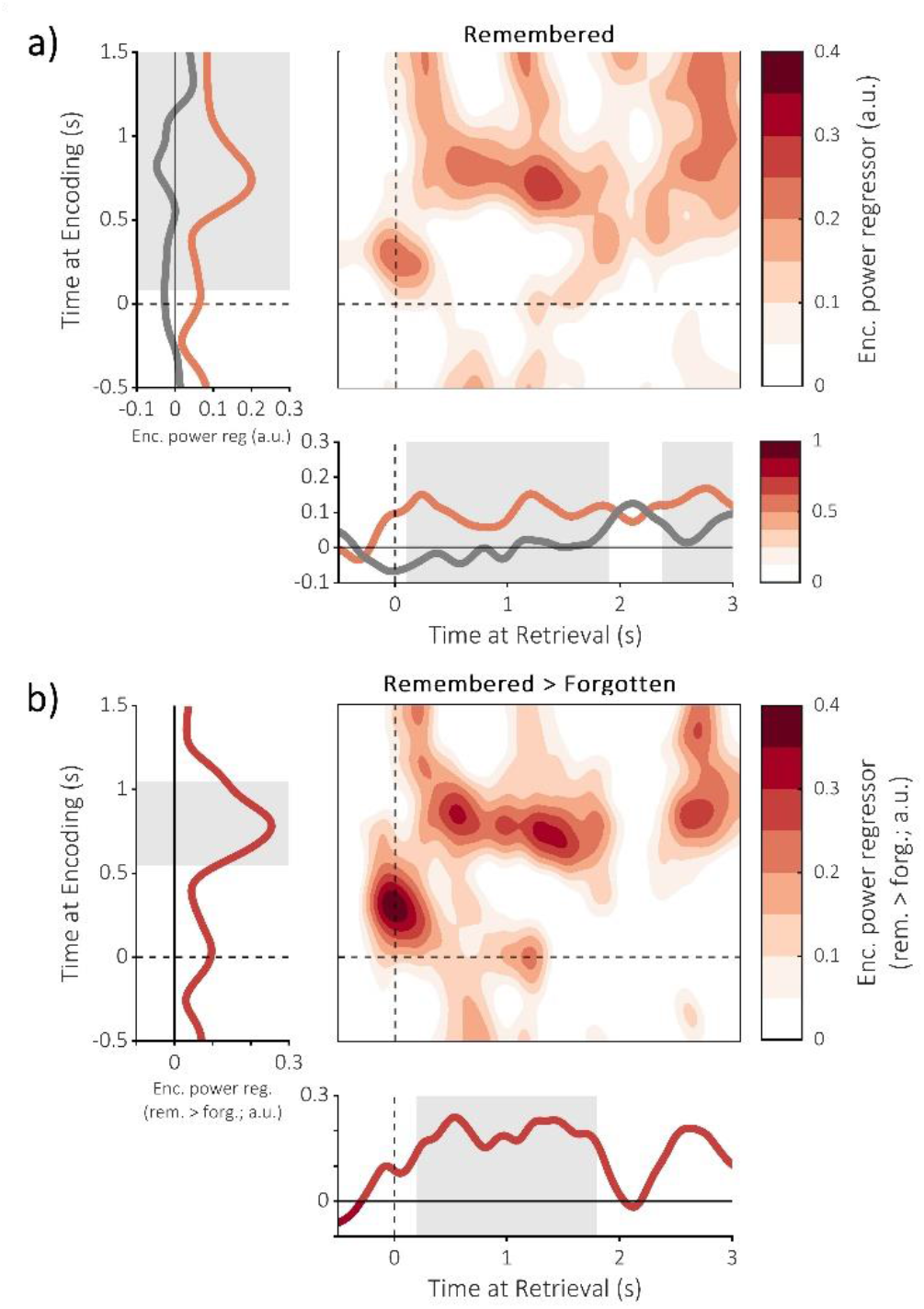
Temporal specificity of the dependency of retrieval power on prior encoding power. **(a)** time-generalisation matrix depicting the extent to which encoding alpha/beta power (7-15Hz; parietooccipital channels in memory-related cluster) predicts later retrieval alpha/beta power for remembered items. Line plot beneath the matrix depicts how encoding power predicts retrieval power for every time point at retrieval (hits in orange, misses in gray; averaged across encoding time-points included in significant cluster). Shaded area depicts time-points included in significant cluster. Line plot left of the matrix depicts how encoding power predicts retrieval power for every time point at encoding (averaged across retrieval time-points included in significant cluster). **(b)** time-generalisation matrix depicting the extent to which encoding alpha/beta power (7-15Hz) predicts later retrieval alpha/beta power for remembered items relative to forgotten items.

No significant cluster was observed for forgotten trials [largest cluster: p_clus_ = 0.408, summed t-statistic = 433.09, cluster size = 214, Cohen’s d_z_ = 0.49].

When contrasting matrices between remembered and forgotten trials, a significant cluster was observed [p_clus_ = 0.041, summed t-statistic = 1592.09, cluster size = 425, Cohen’s d_z_ = 0.91], which included encoding time-points from 550ms to 1050ms and retrieval time-points ranging from 200ms to 1800ms (see figure 3b). This result corroborates the prior findings, suggesting that the alpha/beta power decreases present differing time-courses during encoding and retrieval. Notably, the absence of an effect along the diagonal attenuates concerns that the observed relation between encoding and retrieval is due to perceptual overlap or attention to the stimulus, both of which should exhibit highly similar time-courses during stimulus presentation.

Another cluster can be observed at stimulus onset in both plots presented in figure 3, perhaps reflected a similarity in evoked response between encoding and retrieval. However, the statistics associated with this cluster suggests that the size of this effect was no greater than chance when considering remembered triads alone [p_clus_ = 0.451, summed t-statistic = 373.71, cluster size = 148, Cohen’s d_z_ = 0.61]. The same is true when considering the difference between remembered items and forgotten triads [p_clus_ = 0.293, summed t-statistic = 639.93, cluster size = 218, Cohen’s d_z_ = 0.71].

### Alpha/beta power decreases do not carry stimulus-specific information

Lastly, we asked whether the changes in alpha/beta power directly reflect stimulus-specific information. To this end, linear discriminant analysis (LDA) was used to decode stimulus identity during the presentation of the object (where LDA was used to discriminate animate vs. inanimate objects), feature (polka-dot vs. chequered), and scene (indoor vs. outdoor). When running LDA on broadband amplitude, the subcategories of all three stimulus types could be distinguished to a significantly greater degree than chance [animate vs. inanimate: p_clus_ < 0.001, summed t-statistic = 639.88, cluster size = 161, Cohen’s d_z_ = 0.96; polka-dot vs. chequers: p_clus_ < 0.001, summed t-statistic = 739.92, cluster size = 141, Cohen’s d_z_ = 1.27; indoor vs. outdoor: p_clus_ = 0.002, summed t-statistic = 376.43, cluster size = 103, Cohen’s d_z_ = 0.87]. We then repeated this analysis, using alpha/beta power in place of broadband amplitude. Here, none of the subcategories could be reliably distinguished [animate vs. inanimate p_clus_ = 0.245, summed t-statistic = 66.84, cluster size = 29, Cohen’s d_z_ = 0.56; polka-dot vs. chequers: p_clus_ = 0.205, summed t-statistic = 76.58, cluster size = 30, Cohen’s d_z_ = 0.62; indoor vs. outdoor: p_clus_ = 0.134, summed t-statistic = 90.77, cluster size = 41, Cohen’s d_z_ = 0.54]. Taken with the key findings above, these results suggest that while stimulus category can be decoded based on the recorded signal, alpha/beta power is not the driver of this result (replicating earlier reports; Griffiths, Mayhew, et al., 2019; Ng et al., 2013; Weisz et al., 2020). In other words, topographic patterns of alpha/beta power do not carry stimulus-specific information.

## Discussion

When recalling an episodic memory, information about the encoded event is rapidly reinstated within the cortex. Intuitively, the information that is retrieved must have been encoded previously (Tulving & Thomson, 1973), introducing a contingency between encoding and retrieval processes. Here, we ask whether such a contingency is observable on a neural level. Indeed, we uncovered evidence to suggest that decreases in alpha/beta power during successful memory formation predict the magnitude of alpha/beta power decreases during subsequent retrieval. Given that we found no evidence to suggest that these power decreases code for stimulus-specific representations, it would seem that these decreases reflect a neurophysiological operation that is reinstated during memory retrieval.

Alpha/beta power decreases following cognitive engagement are a ubiquitous phenomenon, transcending tasks (e.g. Hanslmayr et al., 2009; Obleser & Weisz, 2012; Pfurtscheller et al., 1994), stimulus modality (e.g. Crone et al., 1998; Krause et al., 1994; Pfurtscheller et al., 1994) and even species (e.g. Chatila et al., 1992; Haegens et al., 2011; Popov & Szyszka, 2020; Wiest & Nicolelis, 2003). Given the ubiquity, it stands to reason that these power decreases reflect a highly general process. Theories have proposed that these decreases support information representation by suppressing background noise or by increasing entropy within the network (Hanslmayr et al., 2012). Several studies have provided empirical support for these ideas by demonstrating that alpha/beta power decreases parametrically scale with the amount of information encoded as, or retrieved from, an episodic memory (Griffiths et al., 2020; Griffiths, Mayhew, et al., 2019; Karlsson et al., 2020; Martín-Buro et al., 2020). The current results add further support to this idea. The amount of information that can be retrieved from memory is contingent on the amount encoded (Tulving & Thomson, 1973). If alpha/beta power decreases support the representation of information, one would therefore expect to find this contingency reflected in the relationship between encoding-related and retrieval-related alpha/beta power. Indeed, we found exactly this: the magnitude of alpha/beta power decreases during episodic memory retrieval can be predicted by alpha/beta power decreases observed during encoding. These results support the idea that alpha/beta power decreases relate to a process aiding the representation of stimulus-specific information during episodic memory formation and retrieval.

Our analysis of the central effect’s time course help to further contextualise the link between encoding- and retrieval-related alpha/beta power decreases. Here, we found that brief alpha/beta power decreases during encoding predicted extended alpha/beta power decreases during subsequent retrieval. Critically, the timing of these effects match those reported in previous studies which examined memory-related changes in alpha/beta power. The brief alpha/beta power decreases we observe during encoding match the timing of the subsequent memory effect, where encoding-related power decreases arise around 500ms after stimulus onset and last approximately 500ms (e.g. Fell et al., 2008; Griffiths et al., 2016; Hanslmayr et al., 2009). Similarly, the extended alpha/beta power decreases we observed during retrieval match the timing of the retrieval success effect, including both the rapid-onset power decreases (Waldhauser et al., 2016) and those which last for extended periods of time (>1000ms; e.g. Karlsson et al., 2020; Martín-Buro et al., 2020; Michelmann et al., 2016). Given this overlap, it seems reasonable to suggest that the observed contingency between encoding- and retrieval-related alpha/beta power decreases directly relates to the same neurophysiological phenomena associated with the subsequent memory and retrieval success effects.

Research into episodic memory retrieval has long focused on the reinstatement* of neural representations (Schreiner & Staudigl, 2020). This spans from early work demonstrating the reinstatement of modality-specific neural patterns (e.g. Nyberg et al., 2000; Wheeler et al., 2000) up to more recent work demonstrating the reinstatement of stimulus-specific neural patterns (e.g. Chen et al., 2016; Linde-Domingo et al., 2019; Staresina et al., 2012). A common interpretation of these findings is that these patterns reflect the neural representation of the memory within the cortex. That is, these patterns code for a recalled stimulus. However, our findings cannot be explained under this interpretation. While we were able to observe the reinstatement of alpha/beta power during retrieval, we were unable to decode stimulus identity within topographic patterns of alpha/beta power (matching earlier reports: Griffiths, Mayhew, et al., 2019; Ng, Logothetis, & Kayser, 2013; Weisz et al., 2020). This disparity is emphasised when one considers that the reinstatement of alpha/beta power during retrieval could be detected at single channels and frequencies using traditional univariate approaches, while links between stimulus identity and alpha/beta power were not forthcoming despite using more advanced multivariate approaches. Therefore, the observed reinstatement of alpha/beta power is unlikely to reflect the reinstatement of stimulus-specific information.

Rather than reflecting the reinstatement of stimulus-specific information, our results fit with the idea of the reinstatement of a neurophysiological phenomenon (e.g., the suppression of neural noise as discussed above) that supports the neural code of a stimulus. Intuitively, a process that supports neural representations of stimuli during encoding would also be capable of supporting neural representations of stimuli during subsequent retrieval. Therefore, when a neural representation is reinstated, the supportive processes could also be expected to be reinstated. Critically, this is not to say that prior studies cited above are measuring the reinstatement of supportive processes – several of these studies use multivariate approaches to identify fine-grained differences in neural activity for highly similar stimuli that cannot easily be attributed to a general, supportive process. Future studies, however, may benefit from acknowledging that both neural representations and the underlying neurophysiological processes can be reinstated during episodic memory retrieval (Tulving & Thomson, 1973) and tailor analysis accordingly.

It is important to note that this conclusion (that is, alpha/beta power is unlikely to reflect the reinstatement of stimulus-specific information) does not generalise to alpha/beta oscillations in their entirety. Indeed, several studies have been able to reliably deduce stimulus identity within the phase component of alpha/beta band activity (Michelmann et al., 2016; Ng et al., 2013; Staudigl et al., 2015). These results are not contradictory with the current findings, as the phase of an oscillation is mathematically independent of the power of said oscillation. As such, it is completely plausible to suggest that the phase of alpha oscillations carries stimulus-specific information, while the power of said oscillation does not.

With evidence to suggest that alpha/beta power fluctuations are reinstated during episodic memory retrieval, it is worth considering whether other neurophysiological phenomena show similar dependencies between encoding and retrieval (or, indeed, encoding and other processes such as consolidation). Hippocampal theta/gamma activity are both intimately tied to episodic memory encoding and retrieval (e.g. Colgin, 2016; Nyhus & Curran, 2010), and therefore have the potential to demonstrate some contingency between encoding and retrieval. For example, the coupling of gamma activity to particular phases of theta is thought to provide a mechanism capable of encoding temporally-organised sequences (Lisman & Idart, 1995; Lisman & Jensen, 2013), and the retrieval of the sequences is presumed to support the retrieval of temporal order. As such, one could anticipate that hippocampal theta-gamma coupling during retrieval is contingent on coupling during encoding, much like that reported here for alpha/beta power. However, evidence also suggests that encoding and retrieval are optimal at different phases of theta (Hasselmo, 2005; Kerrén et al., 2018), and that distinct gamma bands underpin encoding and retrieval (Colgin et al., 2009; Griffiths, Parish, et al., 2019). Given these differences between encoding and retrieval, there is also reason to believe that hippocampal theta/gamma activity does not show the same parametric contingency reported here. In short, while it may be plausible to predict a contingency between encoding and retrieval in other regions and frequency bands, such a dynamic cannot be assumed without further examination.

It is worth considering that changes in spectral power do not directly equate to changes in underlying oscillatory power (Haller et al., 2018), and instead may reflect a change in the underlying 1/f characteristic of the electrophysiological signal (Miller et al., 2009). While several approaches have been developed to separate changes in 1/f-related activity from changes in oscillatory power (Haller et al., 2018; Wen & Liu, 2016), these approaches suffer signal-to-noise issues when used in single-trial based analyses (Griffiths, Mayhew, et al., 2019) such as the regression models used here. However, the absence of a correction method is not a substantial problem in the data presented here. Changes in oscillatory power should present themselves as narrowband peaks in the power spectrum, whereas changes in 1/f-related activity should present themselves as broadband effects that fluctuates as a logarithmic function of frequency. When analysing remembered and forgotten items separately, we observed a combination of these two phenomena (see figure 2b) – both conditions showed a peak in the alpha/low beta ranges, and a tapering effect into the higher beta frequencies. Critically however, the direct contrast of these conditions produced a narrowband peak within the alpha/low-beta band (see figure 2c). Therefore, it would seem reasonable to conclude that the memory-related (i.e. remembered > forgotten) overlap between encoding and retrieval power was linked to narrowband alpha/beta oscillations (matching earlier reports suggesting these power decreases are oscillatory in nature; Fellner et al., 2019), whereas the broadband overlap in power was linked to another phenomenon present in both the remembered and forgotten conditions, such as perceptual overlap between the to-be-encoded stimuli and the retrieval cue.

In conclusion, we find evidence to suggest that alpha/beta power decreases during the retrieval of an episodic memory are contingent on the magnitude of alpha/beta power decreases that arose during the initial encoding of the memory trace. As alpha/beta power does not code for stimulus-specific information, but rather provides favourable conditions for information representation, these results suggest that neurophysiological operations are reinstated during episodic memory retrieval in order to support the representation of information about the retrieved memory.

## Acknowledgements

T.S. is funded by the European Research Council (grant no. 802681). S.H. is funded by the European Research Council (grant no. 647954) and the Economic and Social Research Council (ES/R010072/1). B.S. is funded by the Wellcome Trust (107672/Z/15/Z).

## Footnotes

* As defined by a recent consensus statement (Genzel et al., 2020). Specifically, “reinstatement” refers to the activation of patterns present during encoding at a later time point while the subject is in an awake state.

